# Glial and BBB modifications correlate with early anxio-depressive-like behaviors and cognitive inflexibility in 3xTg-AD mice

**DOI:** 10.64898/2026.03.13.711598

**Authors:** Guillaume Benhora-Chabeaux, Cassandre Morisset, Thibaut Nicod, Damien Mor, Salomé Delabrouille-Cauliez, Lidia Cabeza, Adeline Etievant, Fanchon Bourasset

## Abstract

Multiple lines of evidence indicate that alterations in glial cells and the blood-brain barrier (BBB) contribute to anxiety- and depression-like behaviors in murine models of depression and chronic stress. Although behavioral and psychological symptoms of dementia (BPSD) represent a major feature of Alzheimer’s disease (AD), this relationship has received limited attention in this pathology. Using the 3xTg-AD mouse model of AD at an early pathological stage, this study explored the relationship between BPSD and variations of BBB and glial cell markers in specific brain regions (hippocampus, basolateral amygdala [BLA], and prefrontal cortex). Memory and emotional behaviors were assessed using a battery of behavioral tests. Endothelial tight junction (TJ) proteins, along with astrocyte and microglial markers, were quantified by western blotting or/and immunohistochemistry in the hippocampus, BLA, and prefrontal cortex. While spatial and recognition memory remained intact, 3xTg-AD mice exhibited an anxio-depressive-like phenotype, impaired coping strategies, and reduced cognitive flexibility. Compared with control mice, 3xTg-AD mice displayed an increased expression of TJ proteins in the hippocampus and BLA, increased microglial cell density in the BLA and the dentate gyrus, and fewer and shorter microglial cell branches in the BLA. A principal component analysis revealed a positive correlation between anxio-depressive-like behaviors and altered microglial morphology in the BLA, whereas impaired cognitive flexibility positively correlates with ZO-1 expression and microglial cell density in the hippocampus. These findings demonstrate an early association between the BBB, glial cells and AD-related BPSD symptoms in 3-month-old 3xTg-AD mice.

## Introduction

Nearly 416 million people worldwide exhibit psychological, cognitive and/or pathophysiological dysfunctions related to Alzheimer’s disease (AD) (1), rendering this neurodegenerative disorder the leading global cause of dementia (2). AD has historically been characterized as a condition in which memory loss arises from progressive neuronal and synaptic degeneration (3), with the two main hallmarks being β-amyloid (Aβ) aggregation and hyperphosphorylated Tau protein (pTau) (4). The last decades, a novel paradigm emerged incorporating anxiety, depression and other emotional alterations into the symptomatology of the disease, grouped under the terms “behavioral and psychological symptoms of dementia” (BPSD) (5), which may affect up to 98% of AD patients (6). From a pathophysiological perspective, AD is also characterized by variations of the neurogliovascular unit (NGVU), including the blood-brain barrier (BBB) and the glial cells (7,8), that can emerge early in the progression of the disease (9,10). Several *in vivo* studies using dynamic contrast-enhanced magnetic resonance imaging (11–13), and *post-mortem* tissue analyses (7,14,15), have reported increased BBB permeability in AD patients. In contrast, other studies have not detected BBB disruption (16–18). Thus, BBB dysfunction in AD patients remains inconsistent and seems to depend on multiple factors, including the chosen experimental methodology, the stage of the disease and the associated comorbidities. The BBB has also been the subject of numerous preclinical studies, using mouse models of AD able to reproduce one or more aspects of the disease, such as cerebral accumulation of Aβ (Tg2576) (19–26), pTau (rTg4510, p25, P301L and P301S) (19,27,28), Aβ and pTau (3xTg-AD) (20,29–32), or *APOEε4* (19,33). The results obtained from these transgenic models are inconsistent, highlighting a need for more studies investigating the BBB in mouse models of AD. Importantly, most of the studies examined the BBB in late stage of pathology, missing alterations that could arise precociously.

Moreover, studies have reported BPSD-like phenotypes in mice models of AD, including 3xTg-AD mice, at various ages and disease stages (for a review, see (34)). Anxious and depressive-like behaviors have been recently reported in 2- and 4-month-old 3xTg-AD mice (35,36), suggesting that neuropsychiatric alterations may precede major cognitive decline in this model. However, the underlying neurobiological mechanisms of these anxious and depressive-like manifestations remain insufficiently characterized, and the literature is fragmented regarding the precise nature and onset of this symptomatology.

Recent evidence suggests that glial cells and BBB are key neurobiological mediators of anxious and depressive symptomatology in patients with neuropsychiatric disorders, and in mouse models of stress and schizophrenia (37–42). In this line, we recently showed that decrease expression of the tight junction (TJ) protein ZO-1 correlates to increased BBB permeability in the ventral tegmental area in a mouse model of chronic stress, which might be an underlying driver to motivational deficits (43). Building on observations connecting behavioral alterations to BBB and NGVU variations in mice models of stress, we hypothesized that BBB and glial modifications may also contribute to emotional disturbances in mouse models of AD.

The aim of this study was therefore to explore both BPSD and BBB/NGVU integrity in the 3xTg-AD mouse model at an early stage of the pathology, before the onset of the amyloid plaques and tau tangles (3 months) (44). This study aimed thus at determining whether BPSD could be linked to BBB/NGVU modifications, through (1) a systematic behavioral characterization, (2) an evaluation of BBB/NGVU structural and functional components, and (3) a principal component analysis (PCA) to explore their potential association.

## Materials and methods

### Animals

All animal procedures were approved by the local Ethical Committee in Animal Experimentation from Besançon (CEBEA-58) and authorized by the Ministry of Education and Research. All efforts were made to minimize animal suffering during the experiments according to the Directive from the European Council of the 22nd of September 2010 (2010/63/EU).

A total of 70 animals (35 wild type –WT, and 35 3xTg-AD, males and females 50/50 %) were used in this study. 3xTg-AD (purchased from *Jax*® and bred in our animal facility (B6;129-Psen1tm1Mpm Tg; APPSwe, tauP301L)1Lfa/Mmjax) and WT mice (C57BL/6JRJ, *EtsJanvier Labs*, Saint-Berthevin, France) were group-housed (two to six per cage) under standard environmental conditions (12 h light/dark cycle; temperature: 22±2 °C; humidity: 55±10%) with *ad libitum* access to filtered water and standard chow (KlibaNafag3430PMS10, *Serlab*, Kaiserau, Switzerland).

### Experimental design

From 2 months of age (8 weeks), mice were gently handled (45), and weighed once per week until euthanasia. At 3 months of age (12 weeks), they were assigned to the following groups: (i) a first group to evaluate recognition and spatial memory, as well as BPSD-like phenotype using a battery of behavioral tests (see 2.3 Behavioral experiment; n=20 WT, n=20 3xTg-AD); animals were euthanized and brains collected for immunofluorescence and western blot at 3.5 months of age; (ii) a second group was tested in an operant learning paradigm (n=10 WT, n=10 3xTg-AD); and (iii) a third group to evaluate BBB permeability via dextran diffusion analysis (n=5 WT, n=5 3xTg-AD).

### Behavioural experiment

All the behavioral tests were performed during the light phase (between 8 a.m. and 3 p.m) to minimize circadian influences, and only one test was conducted per day. Mice were transferred to a noise-attenuated experimental room at least 30 minutes before testing. All testing apparatus were cleaned with 70° ethanol between trials and between animals.

#### Light Dark Box (LDB) test

The LDB test was used to assess anxious-like behavior in mice. The apparatus consisted of a 35 cm^2^ opaque box placed on an infrared-reflective platform. The box was divided into two compartments: a brightly lit (∼150 lux) anxiogenic compartment, and a dark (∼10 lux) one covered with red acrylic glass, allowing infrared tracking in dim light. A small opening connected both compartments. To start, mice were placed individually in the dark compartment, and their exploratory behavior (i.e., time spent and number of entries in the lit compartment and total distance traveled) was recorded during 10 minutes with *Viewpoint 3.10* software (*ViewPoint Behavior Technology*, Lyon, France). The latency to the first entry into the lit compartment was measured manually.

#### Open Field (OF)

The OF test was used to further evaluate anxious-like behavior in mice (46). The arena consisted of a square opaque white Plexiglas (L 50 cm x l 50 cm x h 24 cm), brightly lit in the center (∼150 lux) and virtually divided into a peripheral zone and a central zone (20 cm x 20 cm). Mice were individually placed in the central zone and allowed to explore the apparatus for 10 minutes. Behavior was recorded with *Viewpoint 3.10* and the total distance traveled, the immobility time and the time spent in the central zone were measured.

#### Novel Object Recognition (NOR) test

The NOR test assesses recognition memory (47) and it was conducted in the OF arena, 24h later and under attenuated lighting (∼40 lux). In a first phase of familiarization, two identical objects (transparent ridged glasses or smooth plastic pepper shaker) were placed diagonally (15 cm from walls) and the animals were left 10 minutes to freely explore. In the second phase of testing, 3 hours later, one familiar object was replaced with a novel object, respecting the position of the diagonal in the arena. The familiar and novel objects and their positions were randomly placed during trials. Exploration time of the familiar and novel objects were manually scored with *BORIS* software (*Behavioral Observation Research Interactive Software*, Torino, Italy) (48). Trials lasted at least 5 minutes, extended until 20 seconds of cumulative exploration were reached (47). Mice failing this criterion were excluded. The discrimination index was calculated as [(time novel object – time familiar object) / (time novel object + time familiar object)]. A pilot study confirmed no innate object preference (data not shown).

#### Spontaneous alternation in a Y maze

This test assesses spatial working memory (49). The Y-maze consisted of three white opaque Plexiglas arms (35 cm × 20 cm, 125° apart), called A, B and C. Mice were placed at the beginning of arm A and allowed to explore the maze for 8 min. The proportion of alternation ([number of alternations / (total arm entries – 2)] × 100) was then calculated (49).

#### Rotarod testing

Motor coordination was assessed using an automatized rotor with a 3 cm diameter (Mouse RotaRod NG, *Ugo Basile SRL*, Gemonio, Italy). After a first training trial (15 rotation per minute (rpm) for 120 seconds), mice underwent an accelerating trial (10 to 45 rpm over 60 seconds, then constant 45 rpm for 60 seconds). Behavior was video-recorded, and the latency to fall was manually scored.

#### Cliff avoidance reaction (CAR) test

A modified CAR test (50) was used to assess coping strategies in a low-aversive environment. Each mouse was placed on a round transparent platform (an inverted glass beaker d: 18 cm, h: 27 cm) illuminated at 30 lux. Behavior was video recorded until animals jumped or fell, with a maximal duration of 10 minutes.

#### Splash Test (ST)

The ST assesses depressive-like behavior in mice (51). After 1 hour in individual cages, mice received two sprays of 10% sucrose solution (D(+)-Saccharose, *Roth*, Karlsruhe, Germany) on the dorsal coat near the tail base. Animals were recorded for 5 min, and grooming behavior, as an index of self-centered behavior (51), was manually scored using *BORIS*.

#### Forced Swim Test (FST)

The FST assesses mice coping strategies in response to an acute inescapable stress (52), reflecting anxious- or depressive-like behavior (53) and cognitive flexibility (i.e., the ability to switch between strategy) (54). Mice were placed for 6 minutes in a glass cylinder (d: 18cm, h: 27cm) filled with water (27°C ± 1°C, 40 lux). Their behavior was video-recorded, and the swimming time (from 2 to 6 minutes) was manually scored using *BORIS*. Water was changed between animals.

#### Operant conditioning and cognitive flexibility

Prior to testing, mice were food restricted to 80-90% of their baseline weight. The test was performed as previously described (55). After 3 days of habituation (10 min/day) in operant chambers (*Med Associates*, Hertfordshire, UK), mice were trained to nose-poke under a Fixed Ratio (FR)-1 schedule. A food pellet (20 mg Dustless Precision Pellets® Grain-Based Diet, *PHYMEP s.a.r.L.*, Paris, France) was delivered to the container by a single nose-poke in the active hole. Acquisition was defined by ≥3:1 active/inactive response ratio, ≥20 rewards per session (1hour maximum or 50 rewards per session), consistent across 3 sessions. After contingencies acquisition, training shifted to FR-5 (five consecutive nose-pokes in the active hole for a single food pellet delivery) for three consecutive days. The FR-5 training was followed by a reversal learning (RL) task, where contingencies for active/inactive holes were switched. RL testing was performed in four sessions (2/day, 1 hour maximum or 50 rewards per session). Performance was quantified as the percentage of correct responses and individual learning rates.

### Tissue processing

#### Immunofluorescence

Half of the animals from the first group (n=20) were euthanized with pentobarbital (140 mg/kg, i.p., Exagon, *Med’Vet*, France), and transcardially perfused with 0.9% NaCl followed by 4% paraformaldehyde solution (PFA, *Roth®*, Karlsruhe, Germany; in 0.1 M phosphate buffer (PB), pH 7.4). Brains were post-fixed overnight at 4°C, cryoprotected by immersion in 15% sucrose solution (D(+)-Saccharose, *Roth*, Karlsruhe, Germany; in 0.1 M PB) for 24 h, and frozen by immersion in isopentane (2-methylbutane, *Roth*, Karlsruhe, Germany) maintained at -70°C. Brains were sliced in 30-µm-thick coronal sections using a cryo-microtome (Microm KS 34-S: 47 282; cooling system: R404a, *Micron GmbH*, Walldorf, Germany) and stored in cryoprotectant (50% PB, 25% glycerol, 25% ethylene glycol) at –20 °C till being processed by immunofluorescence.

#### Western blot

The remaining animals of the first group (n=20) were euthanized by cervical dislocation under ketamine/xylazine (100/10 mg/kg, Med’Vet, i.p.) anaesthesia. Hippocampi were then rapidly dissected on ice, snap-frozen in liquid nitrogen (–75 °C) and stored at –80 °C until processing by western blotting.

#### Dextran permeability

As we previously described (43), mice were injected via the tail vein with Texas Red–labelled 40 kDa dextran dissolved in distilled water (3 mg/kg; D1829, *Fisher Scientific*) under isoflurane anaesthesia. After 20 minutes, mice were euthanized by cervical dislocation. Brains were removed and fixed by immersion in 4% PFA (24 h, 4°C), cryoprotected in 15% sucrose solution, and frozen in isopentane (–70 °C). Brains were sliced in 30-µm-thick coronal sections using a cryomicrotome (Microm KS 34-S: 47 282; cooling system: R404a, *Micron GmbH*, Walldorf, Germany) and mounted on gelatine-coated slides immediately before immunofluorescence processing (see 2.5 Immunofluorescence staining and imagery).

### Immunofluorescence staining and imagery

Five-to-ten individuals of each experimental group from cohorts 1 and 2 were randomly selected for immunofluorescence approaches. Sections followed an antigen retrieval protocol aiming at maximizing antigenic site exposure for antibody binding (10 mM citrate buffer, pH 6.0, 96 °C, 30 min). After washing with PBS-Triton 0,3%, sections were incubated with primary antibodies: IBA-1 (1:1000, *Abcam*, ab178846, rabbit), GFAP (1:500, *Abcam*, ab7260, rabbit), ZO-1 (1:500, *Santa Cruz,* sc-33725, rat) and/or Tomato-Lectin (1:500, *Sigma-Aldrich*, L0401) diluted in a blocking buffer (milk and bovine serum albumin) for 24 h at room temperature. Sections were then incubated in secondary antibodies solutions, either with Alexa anti-rabbit IgG (1:2000, *Invitrogen*, A-11008, goat) and Cy3 anti-rat IgG (1:500, *Invitrogen*, A10522, goat), overnight at room temperature. Sections were then cover slipped with mounting medium (40% PB, 60% glycerol; *Roth®*, Karlsruhe, Germany).

Photomicrographs of the regions of interest [infralimbic cortex (IL), basolateral amygdala (BLA), dorsal hippocampus subregions -dentate gyrus (GD), Ammon’s horn 1 and 3 (CA-1 and CA3)], chosen based on their involvement in mnesic and emotional processing, were acquired using a ZEISS AxioImager.Z2 microscope, equipped with ApoTome.2 and a ORCA-Flash4.OLT camera (*Zeiss*, Germany). 10x and 20x objectives and identical parameters of exposition time and intensity between slices were used. Cell density, microglial morphology (ramification endpoints, process length) and fluorescence intensity related to IBA-1 and GFAP expression were quantified using *ImageJ/Fiji* (56) software (National Institute of Health, Bethesda, Maryland, USA). ZO-1 and Lectin-FITC occupied surface and fluorescence intensity were also quantified.

For 40 kDa Dextran-permeability assays, 30 µm coronal sections were mounted on gelatin-coated slices and incubated with Tomato-Lectin (1:500, Sigma-Aldrich, L0401) for 24 hours. After washing, slices were observed and pictures of the same regions of interest were taken with the ZEISS Axio Imager.Z2 microscope. Photomicrographs were acquired using 20x Z-stack. Z-stack were then transformed into maximal intensity projection to measure fluorescence with the Zen2 pro (blue edition) version 2.0.0.0 software (©Carl Zeiss Microscopy GmbH, 2011), using ROI spline tool (57) to delineate an area in the vessel dyed with Lectin and around it. Dextran-FITC fluorescence was measured inside and outside of several brain microvessels, in different slices and for each animal. A ratio was used to approximate the extravasation of the dye, by dividing the fluorescence in the outside of vessels by the fluorescence inside the vessels, as previously described by our team (43).

### Western blotting

Hippocampi were lysed in RIPA buffer (50 mM Tris-HCl pH 7.4, 150 mM NaCl, 1 mM EDTA, 1% Nonidet P40, 0.5% sodium desoxycholate) with protease and phosphatase inhibitors (*Roche Diagnostics*, Meylan, France). Lysates were sonicated, centrifuged (10,000× g, 10 min, 4 °C) and supernatants were stored at –80 °C. Protein concentration was measured using a Bio-Rad protein assay (*Bio-Rad,* Marnes-la-Coquette, France). Samples (30 µg) were solved in Laemmli buffer (*Bio-Rad*) and separated by a 4–15% Mini-PROTEAN® TGX Stain-Free™ Protein Gels SDS-PAGE (*Bio-rad*), transferred to PVDF membranes (*Bio-rad*) and visualized with ChemiDoc MP (*Bio-Rad*). Stain-free total protein was used for normalization. Membranes were blocked (5% milk in PBS-T (0.5 mM Tris-HCl, 45 mM NaCl, 0.05% Tween 20, pH 7.4) and incubated with primary antibodies: Claudin-5 (1:1000, *Sigma-Aldrich*, ABT45, rabbit), ZO-1 (1:500, *Santa Cruz,* sc-33725, rat) or occludin (1:200, *Santa-Cruz*, sc-133256, mouse). Bound primary antibodies were detected using fluorescent secondary antibodies StarBright Blue 520 anti-rabbit IgG (1-5000, *Bio-rad*, 12005870, goat), Alexa 555 anti-mouse IgG (1-2000, *Invitrogen*, A-31570, donkey) or Cy3 anti-rat Igg (1-2000, *Invitrogen*, A10522, goat). The bands were quantified by densitometry with Image Lab v6.1, normalized to stain-free total protein.

### Data and statistical analyses

The results are presented as means ± SD. The statistical analyses and figures were performed using GraphPad Prism 10 software (*GraphPad Inc.*, San Diego, United States). The nature of the data sets was verified using Shapiro-Wilk and Levene’s tests to respectively study the data sets’ normality of distribution and the homogeneity of variance. An unpaired two-tailed t-test was used to analyze the results from two sets of data, except when the data did not follow a normal distribution. In this case, an unpaired two-tailed Mann-Whitney test was used. Statistical significance was defined as p < 0.05. A PCA was conducted to explain the variance across measured variables. The analyses aimed to extract key components underlying the association between behavioral outcomes and neurobiological results.

#### Modelling analysis of the cognitive flexibility (Reversal Learning)

During the RL test, the number of correct answers (allowing reward) was plotted against the number of trials completed by WT and 3xTg-AD mice.

In WT mice, each correct response evolved sigmoidally with the number of trials, as does a dose-response curve. Thus, we applied a modelling approach previously published (58), allowing to mathematically describe the progression curves of cognitive flexibility and to estimate objective parameters. In 3xTg-AD mice, three groups of mice were identified. The pattern of the group#1 was similar to that of WT and was therefore fitted as a sigmoid curve. Mice constituting the group#3 show no correct answer and no progression during the RL test, suggesting that they were not able to adapt their behaviour to obtain the rewards. No fitting curve was calculated since no progression curve was observed. Finally, group#2 showed an intermediate performance: mice adapted their behavior to obtain the rewards, but they needed more trials than mice from the group#1 and WT. A sigmoidal model was also applied to fit the data of group#2.

The sigmoidal equation (Eq. 1) used to fit the observed data was a classical Emax sigmoidal equation:

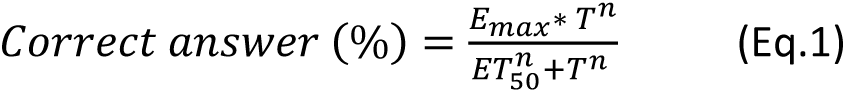

where *T* is the number of trials completed by the mice, *E_max_* is the maximum effect (i.e. correct answers, %) produced by the repetition of the trials, *ET_50_* is the number of trials necessary to reach 50% of the *E_max_*, and n is the sigmoidicity factor. This sigmoidicity factor is an indicator of the curvature of the *E_max_* model.

The fitting was performed for each mouse, using Graphpad Prism ®. Criteria to evaluate the goodness of fit were (i) the visual adequacy between observed and predicted curves and (ii) optimal coefficients of variation of the estimated parameters (CV %).

## Results

### Young 3xTg-AD mice show preserved memory function but exhibit an anxio-depressive–like phenotype

The NOR test was performed to assess mid-term recognition memory in 3-month-old 3xTg-AD mice (Fig. 1a). No difference was observed in the discrimination index between 3xTg-AD and WT mice (t-test: p=0.9262, t=0.09326, df=36), indicating intact recognition memory. Spatial working memory was then evaluated using the spontaneous alternation Y-maze test (Fig. 1b). Similarly, no difference between groups was observed in the percentage of spontaneous alternation (t-test: p=0.1620, t=7.654, df=18), suggesting a preserved spatial working memory (Fig. 1b).

**Fig. 1.**
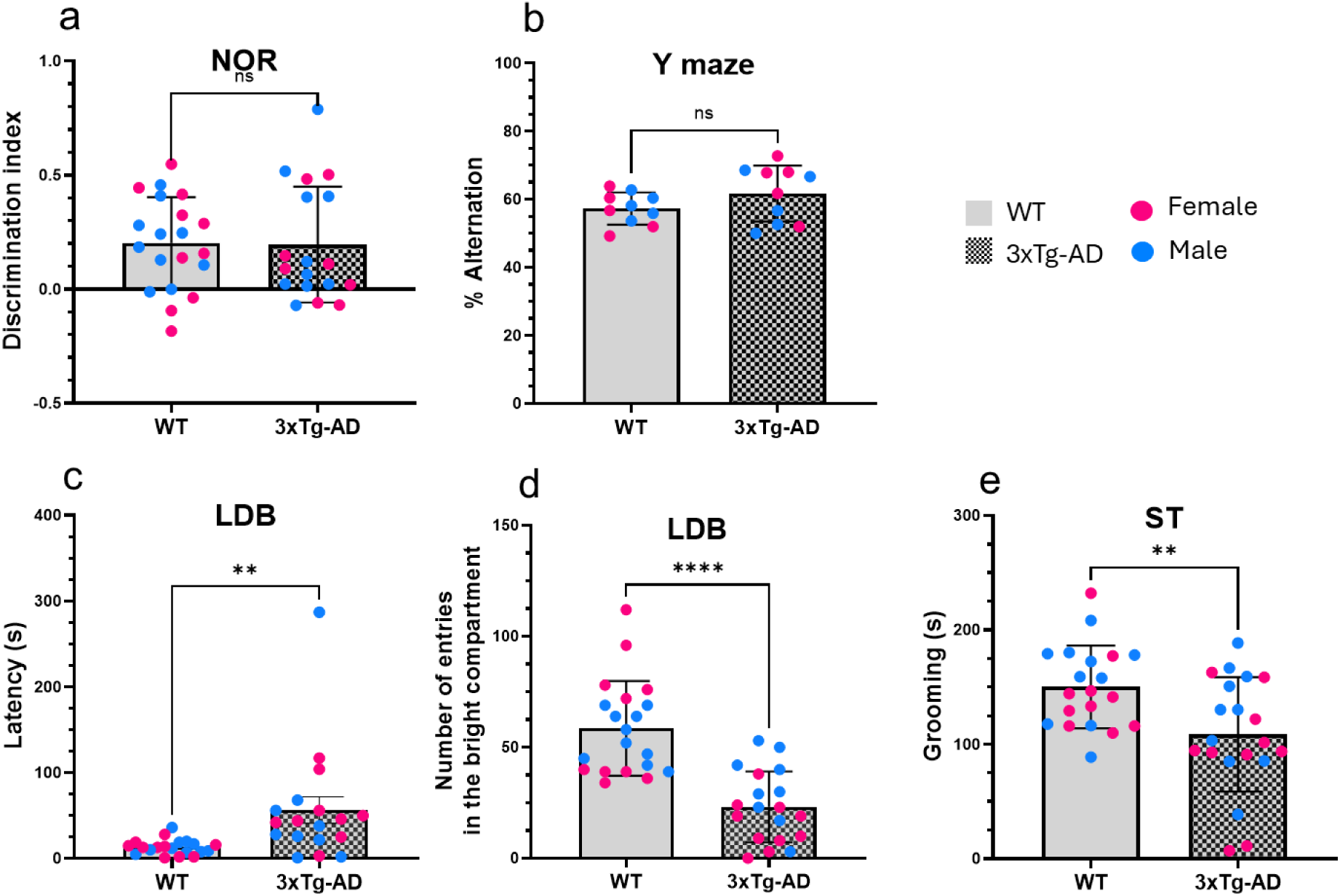
Young 3xTg-AD mice display preserved memory function but exhibit an anxio-depressive–like phenotype. (a) The discrimination index is similar between WT and 3xTg-AD mice in the Novel-Object Recognition test. (b) The percentage of spontaneous alternation is similar between WT and 3xTg-AD mice. (c) The latency of the first step in the bright compartment in the Light-Dark Box test higher for 3xTg-AD mice. (d) The number of entries in it is lower for 3xTg-AD mice. (e) Grooming duration is reduced for 3xTg-AD mice. Blue points represent males and pink points represent females (∼50% of each sex in each mice group). **p<0.01; ****p<0.0001. ns: not significant

Latency to enter in the bright compartment and the frequency of transition into the LDB test are commonly used to assess anxiety-like behaviors in mice (59–61). In this study, 3xTg-AD mice showed a significant increase in the latency to the first entry into the bright compartment (t-test: p=0.0060, t=2.919, df=36) (Fig. 1c) and a marked reduction in the frequency of entries into the bright compartment (t-test: p<0.0001, t=5.846, df=37) (Fig. 1d).

Self-grooming was evaluated using the ST. During the 5-minutes session, 3xTg-AD mice spent less time grooming compared to WT mice (t-test: p=0.0046, t=3.013, df=38) (Fig. 1e). Taken together, these results suggest an anxio-depressive like phenotype in 3 months-old 3xTg-AD mice.

### Decreased exploratory motivation in 3xTg-AD mice

In the LDB test, 3xTg-AD mice showed reduced exploration of the apparatus, reflected by a significant decrease in the total distance travelled (t-test: p<0.0001, t=9.144, df=37) (Fig. 2a). A similar reduction was observed in the OF test (t-test: p<0.0001, t=5.904, df=37) (Fig. 2b). Given that decreased exploration might result from impaired motor coordination (62), we assessed motor function using a Rotarod task. During the simple constant-speed trial (15 RPM), 3xTg-AD and WT mice performed similarly (Mann-Whitney: p=0.4737, U=40) (Fig. 2c), remaining on the rod for the entire 120-seconds trial, indicating preserved motor coordination. In the more demanding ramping-speed paradigm—during which the rotation increased from 10 to 45 RPM followed by a 60-seconds plateau at 45 RPM—3xTg-AD mice outperformed WT controls (Mann-Whitney: p=0.0216, U=22) (Fig.2d).

**Fig. 2.**
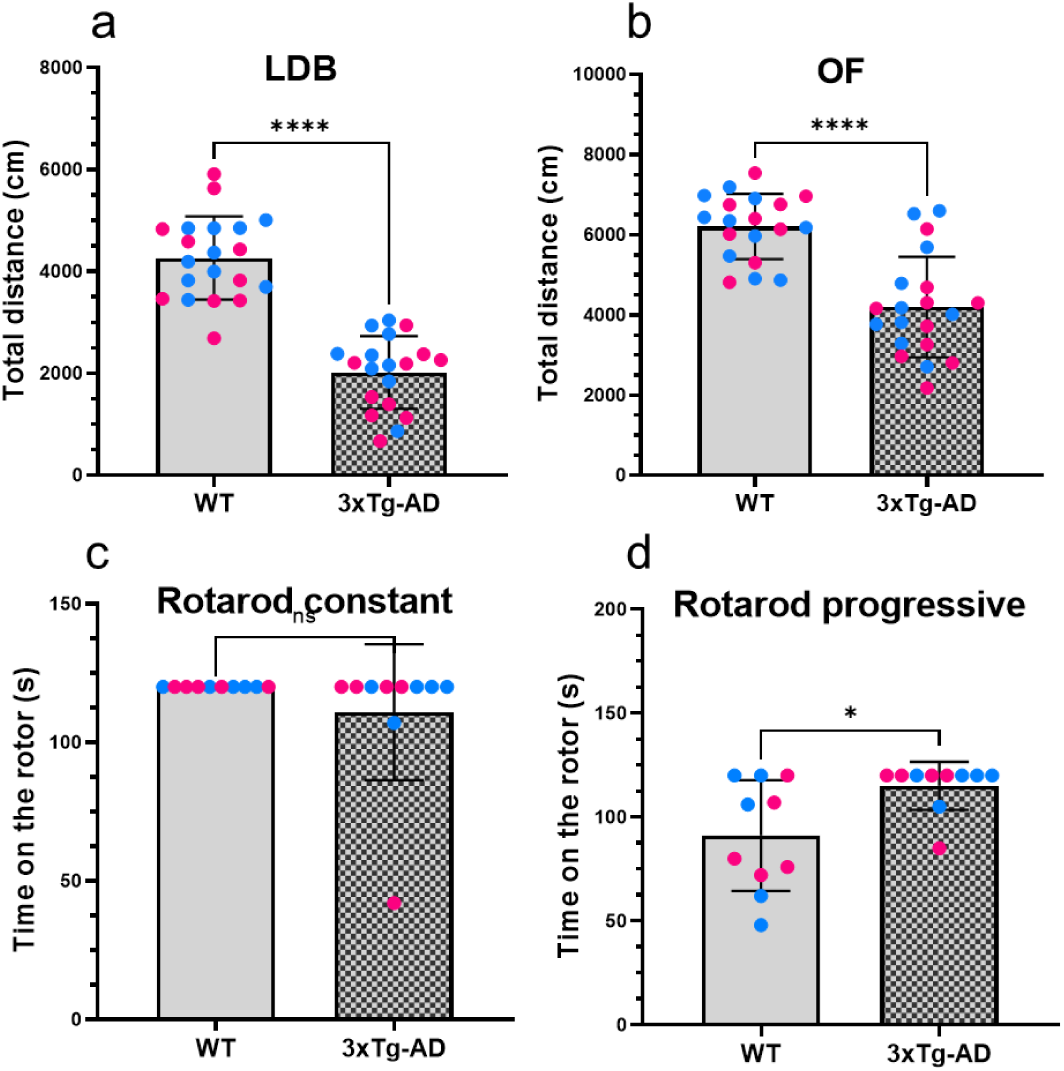
Assessment of exploratory behavior and motor abilities in WT and 3xTg-AD mice. (a) Total distance travelled in the Light-Dark Box test is lower for 3xTg-AD mice. (b) Total distance travelled in the Open Field test is lower for 3xTg-AD mice. (c) Time spent on the rod during the constant-speed Rotarod trial is similar between WT and 3xTg-AD mice. (d) Time spent on the rod during the ramping-speed Rotarod trial is higher for 3xTg-AD mice. Blue points represent males and pink points represent females (∼50% of each sex in each mice group). *p<0.05; ****p<0.0001. ns: not significant

These results indicate that the reduced exploration observed in the LDB and OF tests is unlikely to reflect motor dysfunction. Instead, it may reflect decreased exploratory motivation, potentially driven by elevated anxiety levels in 3-month-old 3xTg-AD mice.

### Inappropriate coping strategies suggest impaired behavioral flexibility in anxiogenic contexts

In the LDB test, no difference was observed concerning the time spent in the bright compartment (Mann-Whitney: p=0.5547, U=168.5) (Fig. 3a). However, the average time spent per entry was significantly higher in 3xTg-AD mice (Mann-Whitney: p<0.0001, U=48) (Fig 3b). Instead of leaving the bright chamber —as WT mice typically do(63,64)— 3xTg–AD mice tended to remain in the center and remain immobile, a pattern compatible with either reduced anxiety and increased risk–taking, or with anxiety–induced freezing.

**Fig. 3.**
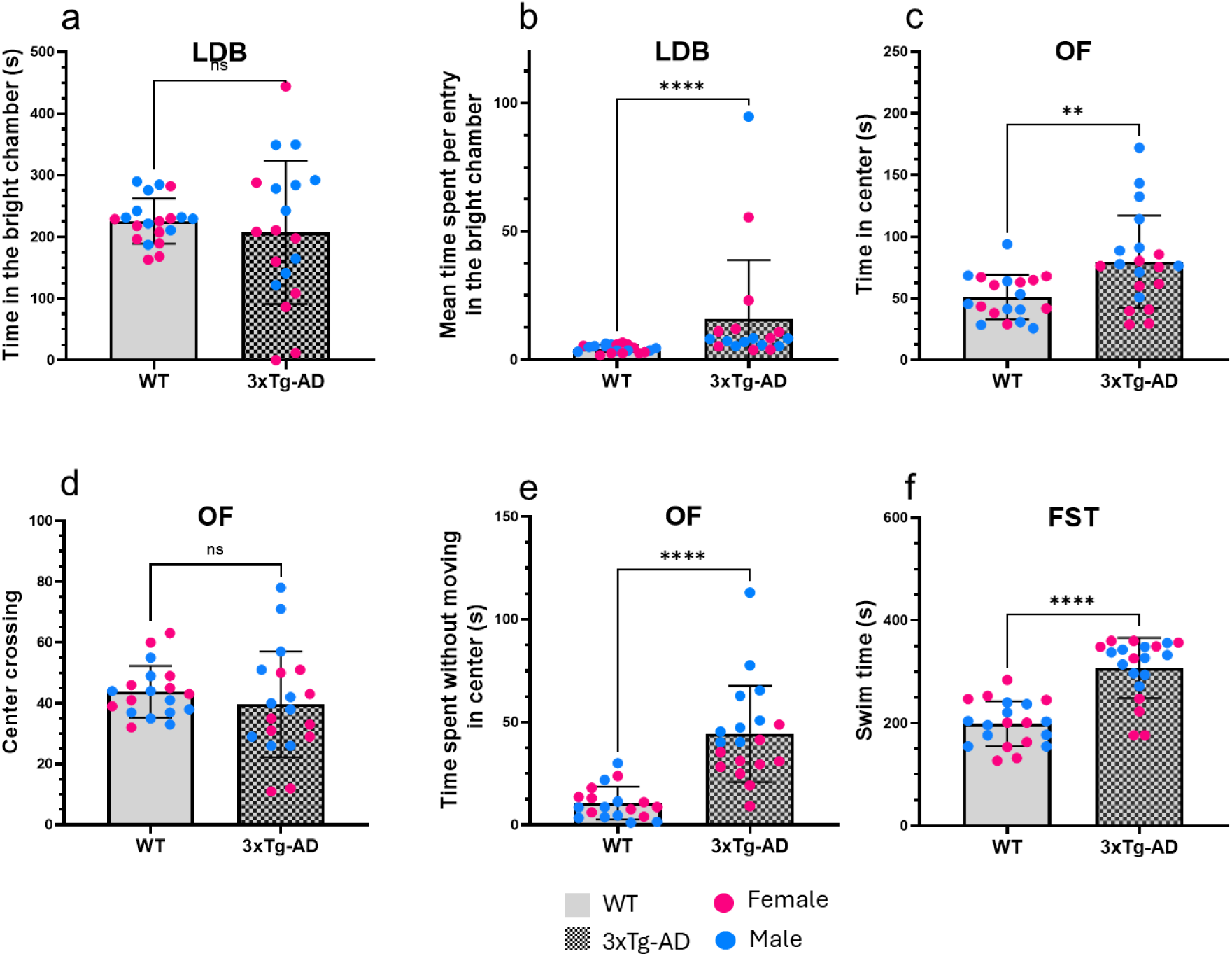
Altered coping strategies and behavioral flexibility in anxiogenic context in 3xTg-AD mice. (a) Total time spent in the bright chamber is similar between WT and 3xTg-AD mice in the Light-Dark Box test. (b) Mean time spent per entry in the bright chamber is superior for 3xTg-AD mice. (c) Time spent in the center of the Open Field test is superior for 3xTg-AD mice. (d) The frequency of crossing the center in the Open Field test is similar between WT and 3xTg-AD mice. (e) Time spent without moving in the center of the Open Field test is superior for 3xTg-AD mice. (f) Time spent swimming in the Forced Swim test is superior for 3xTg-AD mice. Blue points represent males and pink points represent females (∼50% of each sex in each mice group). **p<0.01; ****p<0.0001. ns: not significant

Similar behaviors were observed in the OF test. 3xTg-AD mice spent significantly more time in the centre than WT mice (t-test: p=0.0044, t=3.083, df=37) (Fig. 3c) even if WT and 3xTg-AD show the same number of entries in the centre of OF (t-test: p=0.3627, t=0.9220, df=36) (Fig. 3d). Indeed, 3xTg-AD mice showed an increase in the time spent immobile at the centre of the arena (Mann-Whitney: p<0.0001, U=13) (Fig. 3e).

To differentiate between the aforementioned possibilities, we used a modified version of the CAR test (65) to assess risk–taking in isolation. In this paradigm, 3xTg-AD mice remained significantly longer on the platform than WT mice (see supplemental Figure 1.a), supporting the interpretation of anxiety–driven freezing rather than risk–taking, consistent with an inappropriate coping strategy in aversive situations.

The forced swim test (FST) was then used to examine coping behaviors under acute stress. Rather than interpreting immobility as behavioral despair (66), we followed recent work (52,54,67) suggesting that this test allows to assess behavioral flexibility and the ability to alternate between active and passive strategies when faced with aversive stress. 3xTg-AD mice spent nearly the entire test actively swimming, whereas WT mice alternated efficiently between swimming and immobility, resulting in a significant increase of swimming time in the transgenic group (t-test: p<0.0001, t=6.642, df=38) (Fig. 3f).

Overall, these results indicate that, unlike WT mice, 3xTg-AD mice display abnormally sensitive responses to stressful stimuli and fail to adapt their behavior appropriately.

### Preserved instrumental conditioning but impaired cognitive flexibility in 3-month-old 3xTg-AD mice

To assess their cognitive flexibility, mice were placed in an operant chamber where they had to perform five nose–pokes in a designated correct hole to obtain a reward. All the mice successfully learned to obtain rewards without a significant difference between WT and 3xtg-AD mice (supplemental Figures 1b-c). This training was followed by a reversal learning (RL) task, where contingencies for active/inactive holes were switched, and lasted four sessions.

All WT mice (n= 10) were able to switch holes during the task (Fig. 4a), evidencing their cognitive flexibility. The mean fitting curve is shown in Figure 4b. The mean parameters of the model were: Emax (correct responses, %) = 85.6±8.6 %; ET_50_ = 1.6±0.2 trials and the sigmoidal coefficient, n = 2.8±0.7. These parameters indicate that WT mice reached 85.6±8.6 % of correct responses during the RL and that 50% of the Emax was reached after 1.6 trials.

**Fig. 4.**
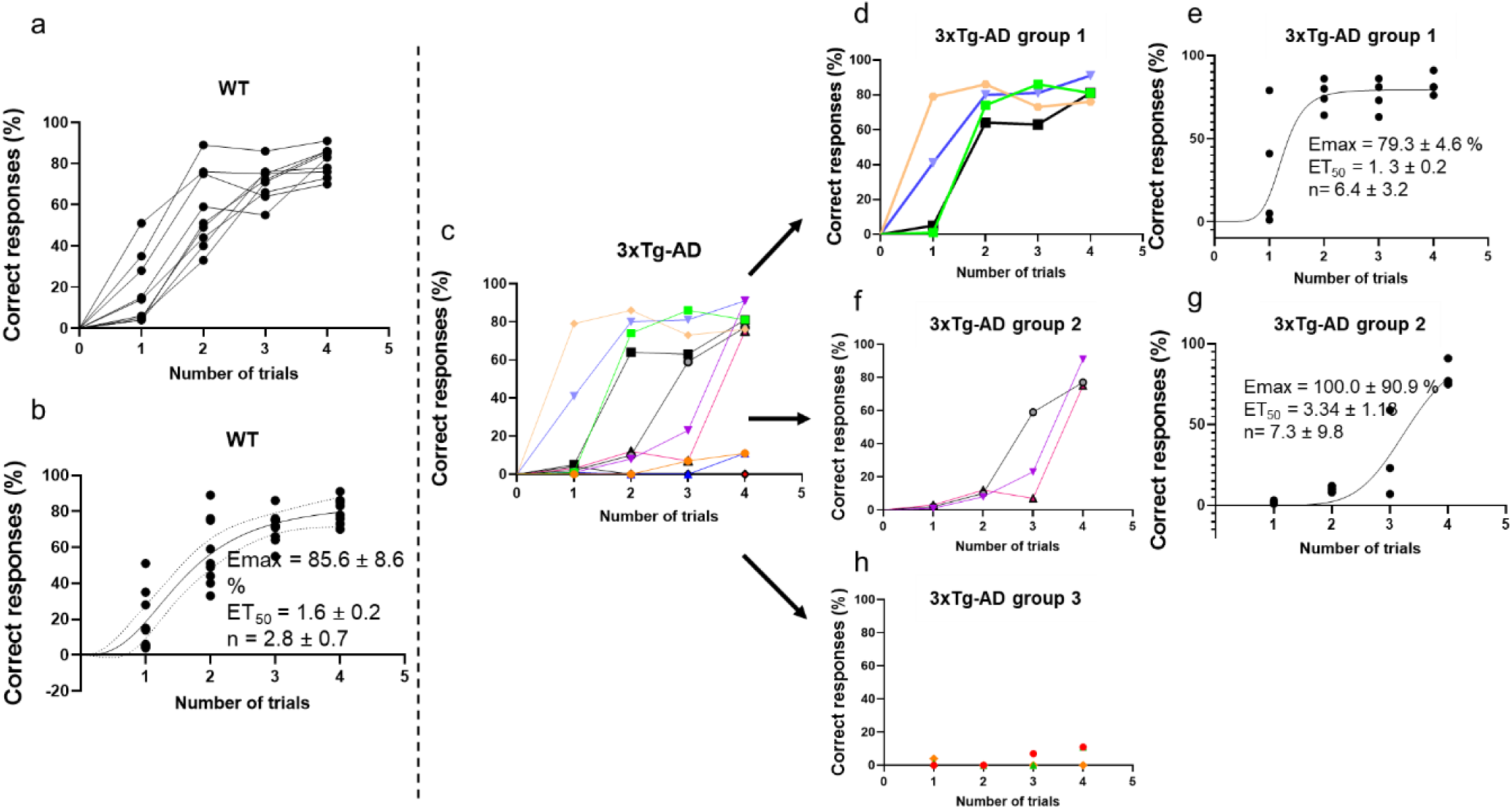
Impaired cognitive flexibility in 3-month-old 3xTg-AD mice. The cognitive flexibility was assessed by the reversal learning (RL) test. Each mouse performed the RL task 4 times. The correct responses in each session were counted. (A-B) The correct responses of each WT mice are plotted against the number of trials. (A) represent the observed data and (B) the observed (points) and predicted (solid line) data obtained by fitting observed data by the sigmoidal equation (Eq. 1). (C) The correct responses of each 3xTg-AD mice are plotted against the number of trials. (D-H) 3xTg-AD mice can be subdivided in 3 subgroups. (D-E) Group 1: (D) the observed correct responses are plotted against the number of trials; (E) the observed (points) and predicted (solid line) data obtained by fitting observed data by the sigmoidal equation (Eq. 1). (F-G) Group 2: (F) the observed correct responses are plotted against the number of trials; (G) the observed (points) and predicted (solid line) data obtained by fitting observed data by the sigmoidal equation (Eq. 1). (H) Group 3: the observed correct responses are plotted against the number of trials (no progression, thus no flexibility, is observed in this subgroup #3)

For 3xTg-AD mice (Fig. 4c), three subgroups of mice were identified. The pattern of the group#1 (n= 4) was similar to that of WT: mice show a progression in the RL test, evidencing the cognitive flexibility of this group#1 (Fig. 4d). Sigmoidal fitting (Fig. 4e) allowed us to estimate the model parameters: Emax =79.3±4.6 %; ET_50_ = 1.3±0.2 trials and the sigmoidal coefficient, n = 6.4±3.2. The Emax and ET_50_ values were very close to those obtained in WT mice, confirming that WT and group#1 of 3xTg-AD mice shared similar flexibility scores.

All mice from the group#2 (n= 3) evolve similarly (Fig.4f) and reached a 100% of correct responses. However, the fitted curve shows that these mice needed more trials to acquire the new contingencies. Hence, ET_50_ = 3.34±1.13 trials, i.e. 3 times that of WT and group#1 of 3xTg-AD mice (Fig. 4g). These mice (group #2) therefore showed an altered cognitive flexibility.

Finally, mice from the group#3 displayed no correct responses and no progression during the RL test, suggesting that they were not able to behaviorally adapt to obtain the reward (Fig. 4h). No fitting curve could be calculated since no progression was observed. These mice (group #3) therefore showed no cognitive flexibility in our experimental conditions.

### Region-specifics laterations in astrocytes and microglia

We performed immunofluorescence analyses of GFAP-labelled astrocytes (density and fluorescence intensity [FI]) (Fig. 5a-c) in the DG, CA1 and CA3, and IBA-1-labelled microglia (density, FI and morphological features) in the DG, CA1, CA3, BLA and IL (Fig. 5d-h) (see supplementary Figure 2 for a detail of all ROI investigated).

**Fig. 5.**
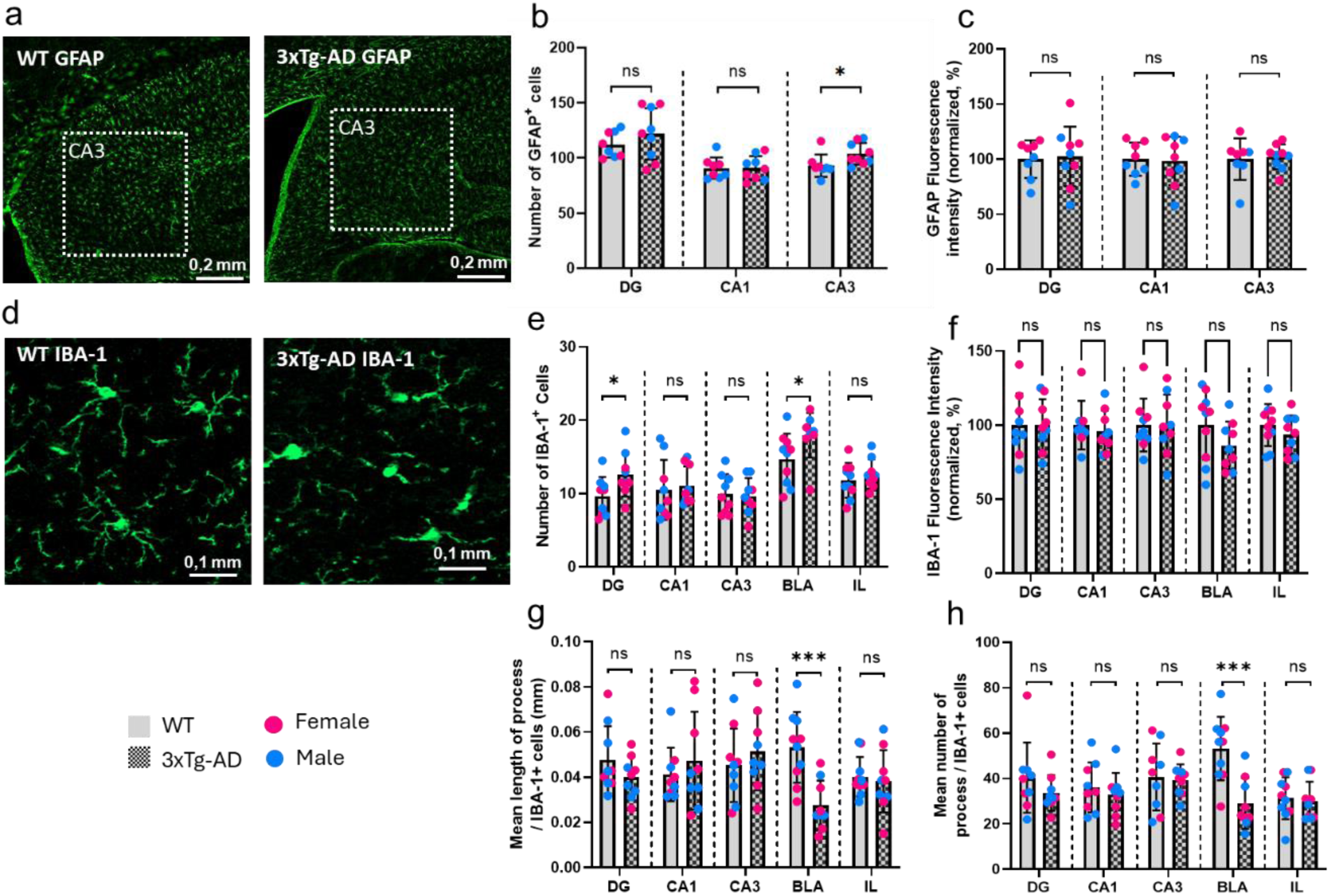
Study of astrocyte and microglial cells. (a) Representative microphotographs of astrocytes labelled with GFAP antibody in the hippocampus of WT (top) and 3xTg-AD (bottom) mice, showing the ROI used for the CA3 subregion. (b, c) Number of GFAP-positive cells per field and fluorescence intensity of GFAP-labeled astrocytes in the Dentate Gyrus, CA1, and CA3 subregions of the hippocampus. (d) Representative microphotographs of microglia labeled with IBA-1 antibody in the Basolateral Amygdala of WT (top) and 3xTg-AD (bottom) mice. (e, f) Number of IBA-1 positive cells and fluorescence intensity of IBA-1-labeled microglia in the Dentate Gyrus, CA1, and CA3 subregions of the hippocampus, as well as in the Basolateral Amygdala and Infralimbic Cortex. (g, h) Morphological features of IBA-1-labeled cells, including the mean process length and the mean number of processes per cell. Blue points represent males and pink points represent females (∼50% of each sex in each mice group). DG: Dentate Gyrus, BLA: Basolateral Amygdala, IL: Infralimbic Cortex. *p<0.05; *** p<0.001. ns: not significant

Within the hippocampus (HPC), the density of GFAP-labelled astrocytes was similar between WT and 3xTg-AD mice in the DG and CA1 region (t-test: p=0.2990, t=1.076, df=15, and p=0.9956, t=0.005572, df=15 respectively), but was significantly higher in the CA3 of 3xTg-AD mice (t-test: p=0.0439, t=2.200, df=15) (Fig. 5b). These observations suggest a proliferation of astrocytes in CA3. FI of GFAP was similar between WT and 3xTg-AD mice in the DG (t-test: p=0.8289, t=0.2199, df=15), CA1 (t-test: p=0.8619, t=0.1770, df=15) and CA3 (t-test: p=0.7816, t=0.2822, df=15) (Fig.5c).

The number of IBA-1 positive cells was significantly increased in the DG and BLA of 3xTg-AD compared to WT mice (t-test: p=0.0421, t=2.209, df=16, and p=0.0459, t=2.153, df=17) (Fig. 5e). However, the number of IBA-1-positive cells were not statistically different between WT and 3xTg-AD mice in CA1 (t-test: p=0.7366, t=0.3422, df=16), CA3 (t-test: p=0.8216, t=0.2292, df=16) and IL (t-test: p=0.4041, t=0.8557, df=17). Similar FI were observed in the regions investigated (t-test: DG: p=0.9829, t=0.02182, df=16; CA1: p=0.5740, t=0.5748, df=15; CA3: p=0.9998, t=0.0002956, df=16; BLA: p=0.1582, t=1.476, df=17; IL: p=0.3146, t=1.036, df=17) (Fig. 5f). Microglia of 3xTg-AD mice had a significant reduced number of processes as well as a reduced average length of processes in the BLA, compared to WT (t-test: number of processes in BLA: p=0.0007, t=4.137, df=17; length of processes in BLA: p=0.0007, t=4.106, df=17) (Fig. 5g-h). On the contrary, no difference was found in the HPC and the IL in terms of length of process (t-test: DG: p=0.2060, t=1.318, df=16; CA1: p=0.4722, t=0.7364, df=16; CA3: p=0.4528, t=0.7696, df=16; IL: p=0.6992, t=0.3930, df=17) and mean number of process (t-test: DG: p=0.2653, t=1.154, df=16; CA1: p=0.5221, t=0.6545, df=16; CA3: p=0.8068, t=0.2486, df=16; IL: p=0.7359, t=0.3428, df=17) (Fig. 5g-h).

### Increased expression of tight junction proteins in specific brain regions of 3xTg-AD mice

To investigate the potential modification of TJ proteins’ expression, western blot and immunofluorescence analyses were performed in several brain regions, in WT and 3xTg-AD mice.

The expression of occludin quantified by western blot in the entire HPC was significantly increased in 3xTg-AD mice compared to WT (Mann-Whitney: p=0.0159, U=1) (Fig. 6a), whereas Claudin-5 and ZO-1 expression remained unaltered (Mann-Whitney: p=0.7361, U=45, and p=0.9682, U=44, respectively) (Fig.6b-c).

**Fig. 6.**
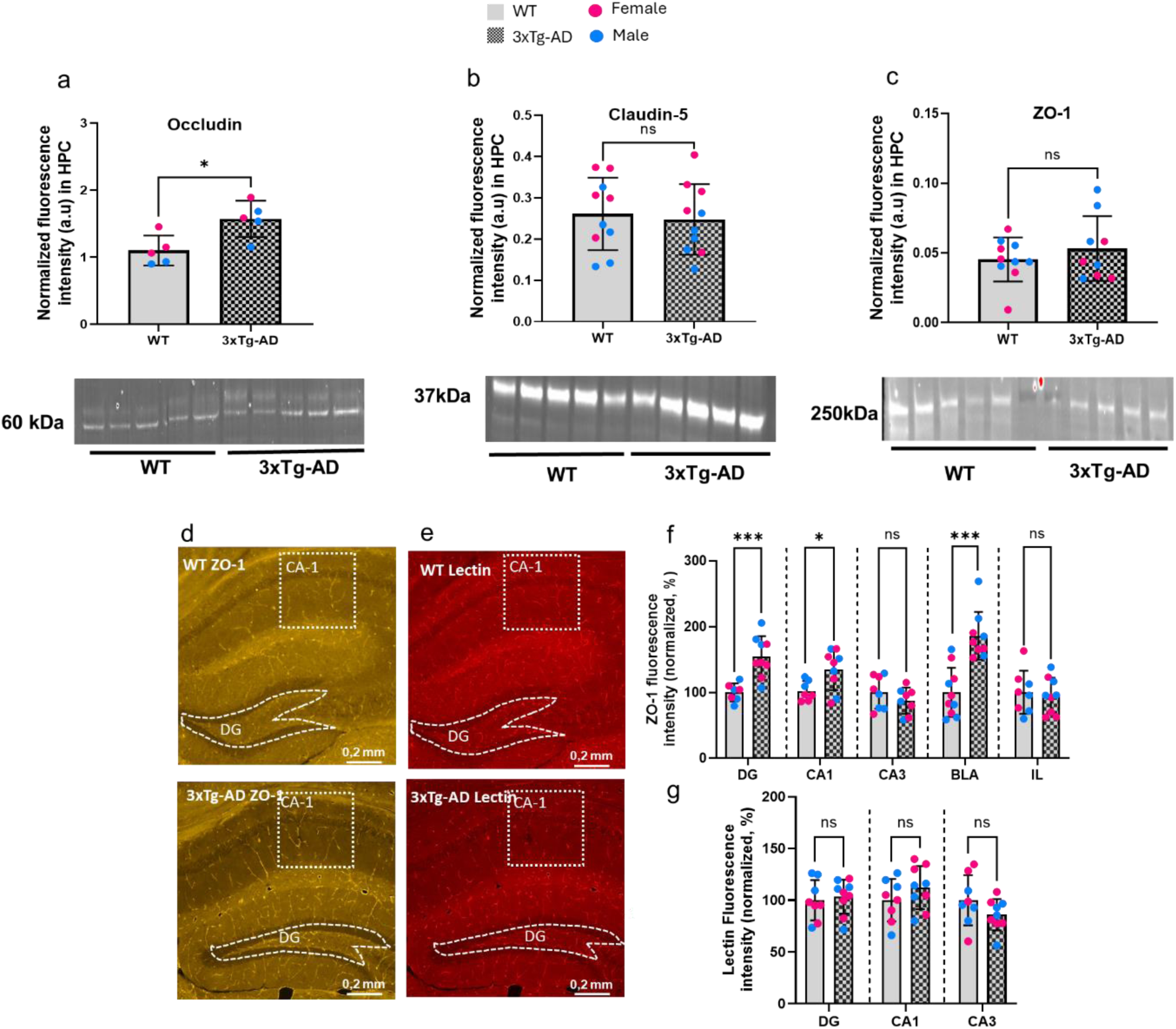
Quantification of lectin and tight junction proteins (ZO-1, Claudin, Occludin) of the BBB through western blot and immunofluorescence. (a, b, c) Analysis of fluorescence intensity standardized to whole protein from western blot of Occludin, Claudin-5 and ZO-1 in the hippocampus of WT and 3xTg-AD mice. (d) Representative microphotographs of ZO-1 in the hippocampus of WT (top) and 3xTg-AD (bottom) mice, showing regions of interest for CA-1 and the dentate gyrus. (e) Representative microphotographs of brains microvessels labelled with tomato-lectin in the hippocampus of WT (left) and 3xTg-AD (right) mice with regions of interest for CA-1 and the dentate gyrus. (f) Normalized fluorescence intensity of ZO-1 in subregions of the hippocampus, the basolateral amygdala, and the infralimbic cortex. (g) Normalized fluorescence intensity of lectin in dentate gyrus, CA1 and CA3 subregions of the hippocampus. HPC: Hippocampus, DG: Dentate Gyrus, BLA: Basolateral Amygdala, IL: Infralimbic Cortex. Blue points represent males and pink points represent females (50% of each sex in each mice group). *p<0.05; ***p<0.001; ns: not significant

To more precisely analyze ZO–1 expression across hippocampal subregions, we measured its FI in 3 subregions: DG, CA1 and CA3. Our results show that ZO-1 FI was comparable between genotypes in CA3 (t-test: p=0.2803, t=1.123, df=14), but was 1.6-fold and 1.3-fold significantly increased in the DG and CA1, respectively, in 3xTg-AD mice compared to WT (t-test: DG: p=0.0007, t=4.343, df=14, and CA1: p=0.0241, t=2.529, df=14) (Fig 6d-6f). To determine whether the increase in ZO–1 could be linked to vessel proliferation, we measured the FI of the blood vessel marker lectin. Lectin-related immunofluorescence was similar between WT and 3xTg-AD mice in DG, CA-1 and CA-3 (p=0.7071, t=0.3831, df=15; p=0.2457, t=1.208, df=15; and p=0.1704, t=1.440, df=15, respectively) (Fig. 6e-6g), suggesting that the increase in ZO-1 expression observed in the DG and CA1 of 3xTg-AD mice occurs independently of blood vessel modifications in these regions. To further study the variation of ZO-1 expression in other brain regions involved in anxious-like behavior and cognitive flexibility, we measured ZO-1 FI in the IL and the BLA. Our results show that ZO-1 FI was 1.9-fold significantly higher in 3xTg-AD mice compared to WT (t-test: BLA: p<0.0001, t=5.078, df=17) but was unaltered in the IL (t-test: IL: p=0.7896, t=0.2711, df=17) (Fig. 6f).

### BBB permeability to 40 kDa dextran is unaltered in 3xTg-AD mice

To study BBB permeability, we measured the FI ratio of FITC-Dextran between the outside and inside of cerebral microvessels, which we also labelled with tomato lectin(43) (Fig 7.a). This ratio allows to quantify potential extravasation of the dye (43), with values superior to 1 indicating BBB leakage. Overall (i.e. DG, CA3, BLA and IL), no difference of BBB permeability was found between WT and 3xTg-AD mice (Mann-Whitney test, p=0,1429, U=2; p=0,9048, U=8; p>0,9999, U=10; p=0.2571, U=6, for DG, CA3, BLA and IL, respectively) (Fig 7.b).

**Fig. 7.**
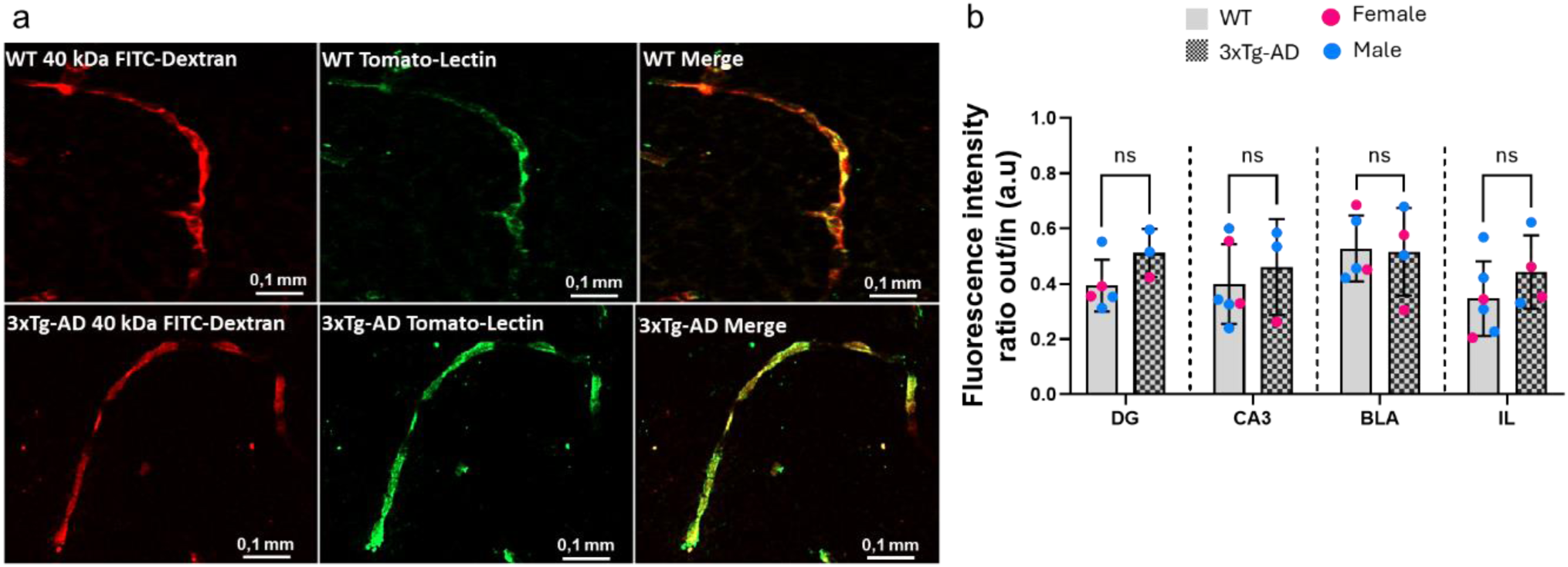
Study of BBB permeability to 40 kDa FITC-Dextran. (a) Representative 20X microphotographs of brain microvessels labeled with 40 kDa FITC-Dextran, Tomato-Lectin, and merged images from WT (top) and 3xTg-AD (bottom) mice. (b) Associated mean values of the outside/inside ratio of Dextran fluorescence in several brain vessels in the dentate gyrus, CA3, basolateral amygdala, and infralimbic cortex. Blue points represent males and pink points represent females (∼50% of each sex in each mice group). DG: Dentate Gyrus, BLA: Basolateral Amygdala, IL: Infralimbic Cortex. ns: not significant

### Principal component analysis suggests BBB and microglia contribution to impaired coping behaviour and anxio-depressive-like phenotype in 3xTg-AD mice

To investigate the relationship between anxio-depressive/flexibility-related behaviors and neurobiological components of the NGVU in 3-month-old 3xTg-AD mice, we performed a PCA using both individual behavioral and neurobiological measurements (n=7 WT, n=7 3xTg-AD). The analysis included the following neurobiological parameters: microglial density (IBA-1+) in DG, CA1 and BLA; GFAP density in DG, CA1; ZO-1 fluorescence in DG, CA1 and BLA; average length of microglial branches in DG, CA1 and BLA) and behavioral parameters (grooming in the ST, latency and time spent per entry in the LDB -LDB-L and LDB-R, respectively, total distance, time in the center and inactivity in the center of the OF, -OF-D, OF-time center, OF IC, respectively-, and swimming duration in the FST).

The PCA identified four principal components (PC1–PC4) explaining over 77% of the total variance, with PC1 and PC2 accounting for ∼56% (Fig. 8a). PC1 separated WT and 3xTg-AD mice into distinct clusters (Fig. 8b). The loading plot and loading values (correlations between variables and PCs) for PC1–PC4, displayed as heatmaps, illustrate the contribution of individual variables to each component (Fig. 8c–d). Inactivity in the center of the OF and ZO-1 IF in the DG loaded strongly and positively on PC1 (loading values: 0.90 and 0.86, respectively) and negatively on OF total distance (−0.75), the average length of microglial branches in the BLA (−0.68), and grooming behavior (−0.56) (Fig. 8c–d). Both grooming and OF measures (OF-D and OF-IC) were primarily associated with ZO-1 expression in the DG and the average length of microglial branches in the BLA (Fig. 8c). FST, OF-time center and LDB measures contributed less to PC1 (loading values: FST: 0.67, LDB-latency: 0.56, LDB ratio: 0.61). FST and OF-time center were associated primarily with hippocampal parameters, including ZO-1 expression in the CA1 (0.76) and microglial density in the DG (0.80). These associations are illustrated in the loading plot (Fig. 8), highlighting variable relationships within PC1 and PC2.

**Fig. 8.**
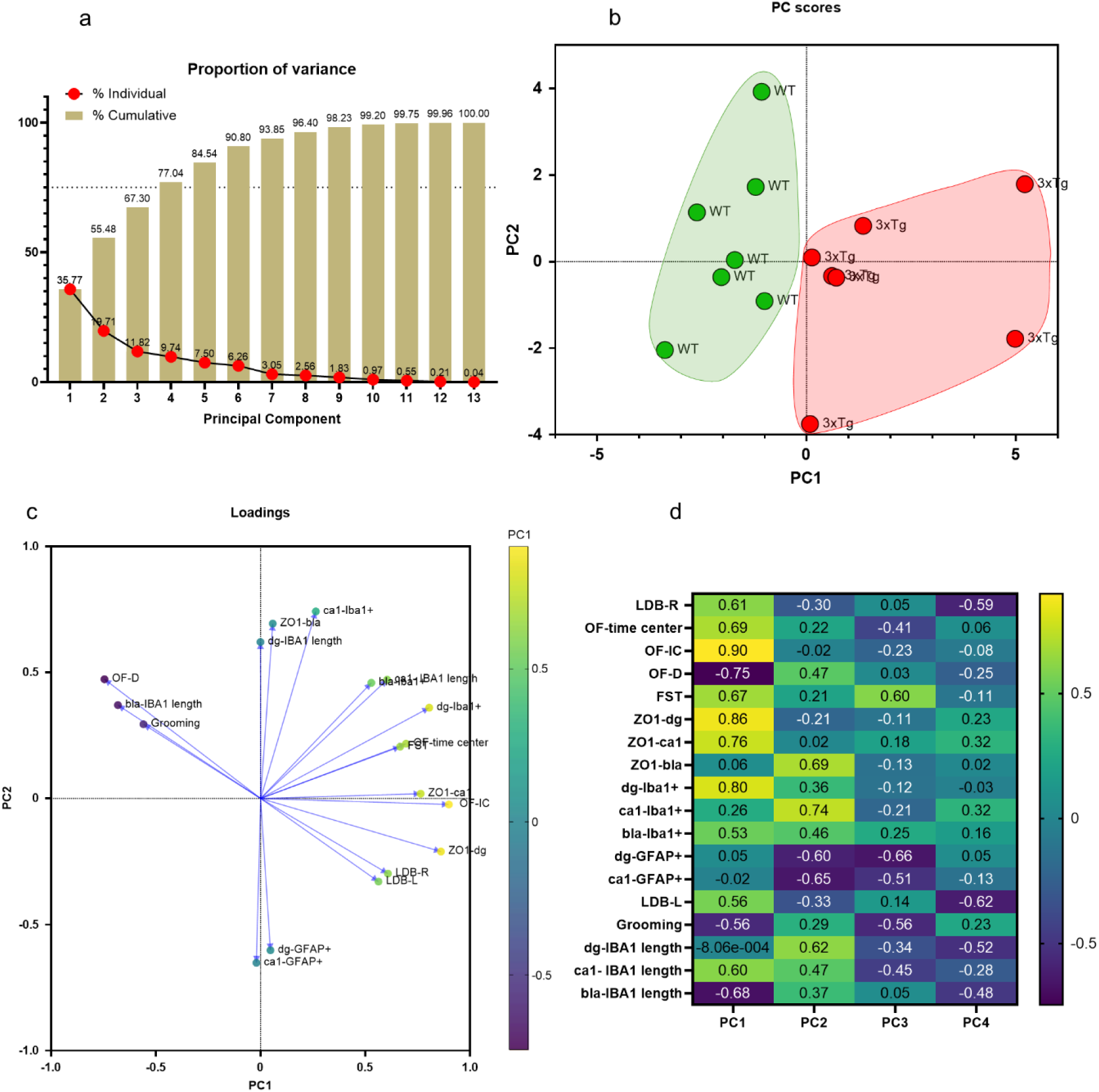
Principal component analysis suggests BBB and microglia contribution to impaired coping behaviour and anxio-depressive-like phenotype in 3xTg-AD mice. Individual behavioral performances of 3xTg-AD and WT mice [Grooming, OF distance (OF-D), time spent in the center of the OF (OF-time center), inactivity in the centre of the OF (OF-IC), LDB latency (LDB-L), time spent per entry in the LDB (ratio: LDB-R) and swimming time in the FST (FST)] were confronted to neurobiological measurements in HPC and BLA [relative fluorescence intensity of ZO-1 (ZO1-dg; ZO1-ca1; ZO1-bla); microglial and astrocyte density (Iba1+ and GFAP+)] through a principal component analysis (PCA). (a) 13 principal components (PC1–13) explain the total variance of the data set, with the four first PCs (PC1–4) explaining 77.04 %. PC1 and PC2 accounting for ∼56%. (b) PC1 separated WT and 3xTg-AD mice into distinct clusters. (c) Loading plot highlights variable relationships within PC1 and PC2 and (d) Loading values for PC1–PC4, displayed as heatmaps, illustrate the contribution of individual variables to each component. Overall, PC1 links behavioral outcomes with variation in ZO-1 fluorescence in DG and CA1, the average length of microglial branches in BLA, and microglial density in DG. ZO-1 in BLA, microglial density in CA1, the average length of microglial branches in DG and GFAP+-cells density in DG and CA1 mainly contributed to PC2

Overall, PC1 links behavioral outcomes with variation in ZO-1 expression in the DG and the CA1, the average length of microglial branches in the BLA, and microglial density in the DG. ZO-1 expression in the BLA, microglial density in the CA1, the average length of microglial branches in the DG and GFAP density in the DG and the CA1 mainly contributed to PC2 (ZO-1-BLA: 0.69; CA1-IBA-1+: 0.74; DG-IBA-1 length :0.62; DG-GFAP+: −0.60; CA1-GFAP+: −0.65).

## Discussion

BPSD, in conjunction with glial and BBB alterations, underscore the multifaceted nature of AD, extending well beyond the classical triad of cognitive deficits and Aβ/Tau deposition. Particularly, modifications of the NGVU, including the BBB, gained increased evidence during the last decades (68–70). However, the potential link between BPSD, BBB and glial modifications has been poorly studied in AD, especially in early stages of the disease. Thus, the objective of this study was to evaluate potential modifications of the NGVU/BBB as well as behavioral abnormalities in the 3xTg-AD mouse model of AD at 3 months of age, i.e., early in the pathophysiological process, before the emergence of distinct memory dysfunction, amyloid plaques, and NFTs.

We found that 3xTg-AD mice display preserved recognition and working memory at this stage, which agrees with previous results obtained in this model (71,72). However, 3xTg-AD mice exhibit anxious-and depressive-like symptomatology, as shown in the ST and LDB test (latency and number of entries in the bright compartment). These findings are consistent with features observed in AD patients, who may present neuropsychiatric symptoms before the onset of marked memory decline, as well as with previous observations obtained in 3xTg-AD mice (35,36,73). Notably, less grooming duration in the ST has been observed in 4 months-old 3xTg-AD mice (35), as well as a decrease of the distance travelled in the OF at 2 and 4 months (35,36). Our results further indicate that 3xTg-AD mice exhibit a generally reduced exploratory behavior in different experimental setups, named the OF and LDB tests (Fig. 2 a-b), suggesting either impaired locomotor ability or abnormal responses to anxiogenic contexts. 3xTg-AD mice outperformed WT mice in the rotarod test, supporting the interpretation that the reduced exploration observed in the OF and LDB reflects abnormal adaptability to anxiogenic situations rather than motor deficits. This interpretation is further supported by the increased time spent in the center of the OF by the 3xTg-AD mice. Our findings are consistent with previous reports indicating that 3xTg–AD mice spend more time in the center of the OF at 6–7 months of age (74–76). These studies interpreted this behavior as either a « disinhibition » (74), a « lower level of anxiety » (75) or an « impaired judgment in assessing danger » (76). By analyzing variables rarely investigated in the OF, we showed that 3xTg-AD mice spent more time immobile in the center of the OF, suggesting the adoption of a freezing-like behavior in response to an anxiogenic environment rather than a disinhibition. Moreover, using the CAR test, we measured risk–taking propensity and found that 3xTg–AD mice, unlike WT, remained on the top of the cliff throughout the entire test, confirming their ability to sense danger, but responding with an atypical, anxiety–driven freezing–like behavior. This reduced adaptability to anxiogenic situations is also illustrated by the results observed in the FST, where 3xTg-AD mice displayed increased swimming times compared with WT controls. This type of behavior has been previously reported in 4, 12 and 17-month old 3xTg-AD mice (35,54,67). Torres-Lista and collaborators interpreted this behavioral response as « a lack of ability of 3xTg-AD mice to shift behavior over time that may result of poorest cognitive flexibility and copying with stress strategies more than behavioral despair *per se*. We show for the first time that this type of behavior is already present in 3-month-old 3xTg-AD mice. Together, these altered behaviors observed in 3-month-old 3xTg-AD mice reflect a reduced ability to appropriately adapt in anxiogenic environments, that might result from impairments in cognitive flexibility, as suggested by Torres-Lista and collaborators (54). This hypothesis is confirmed by results obtained in the RL task, which showed that 60% of 3xTg-AD mice exhibit impaired cognitive flexibility, including 30% that display a complete lack of cognitive flexibility. However, the remaining 40% showed preserved cognitive flexibility, similar to WT mice (Fig. 4). It is noteworthy that our work provides the first evidence that, in addition to anxiety and depression-like symptomatology, 3-month-old 3xTg-AD mice already exhibit a clear alteration of cognitive flexibility.

In order to investigate the hypothesis that these behavioral alterations could be linked to alterations at the NGVU level, we analyzed several glial and endothelial markers, as well as the permeability of the BBB. We show that the density of GFAP-positive astrocytes is slightly increased in the CA3 region of 3xTg-AD mice. Guo and collaborators (77) showed that the area of GFAP-positive astrocytes in the whole HPC was similar in WT and 3xTg-AD aged of 4 months, but was increased at the age of 6 months through F-SMBT-1 imaging (77). Thus, it is possible that studying astrocytes within specific hippocampal subregions allows us to detect alterations that become diluted when analyzing the hippocampus as a whole. A pattern of progressive astrogliosis has also been found in brain of AD patients (78). Importantly, targeting reactive astrocytes exerts a beneficial impact on anxiety in 3xTg–AD mice, suggesting that astrocyte reactivity contributes to emotional disturbances (77,79).

Regarding microglia, ou data show that the density of IBA-1-positive cells is increased in the DG and the BLA of 3xTg-AD mice compared to WT, but remains similar in CA1, CA3 and IL. Our results differ from a previous study that showed no significant difference in microglial density in the hippocampus of 3xTg-AD mice compared to WT mice (80). However, this study did not segment the HPC into subregions as we did, which may contribute to our divergent results. Thus, investigating glial cells like astrocytes and microglia in subregions of the HPC may be useful to bring light on subtle changes that occur during early stages of the disease. Falangola and collaborators found an increase in the density of IBA-1-positive cells in the ventral HPC, subiculum, restrosplenial and cingulate cortex, but not in the dorsal HPC, in 2 months-old 3xTg-AD mice (81), suggesting that microglia proliferation begins in the ventral HPC and could then spread to the dorsal one. A careful examination of the literature fails to identify studies that investigated microglia in the BLA. Our research highlights for the first time that microglial proliferation occurs in the BLA of 3xTg-AD mice at an early stage of the pathology. In addition, we showed a significant decrease of the number of processes per microglial cell as well as a significant decrease of the average length of microglial branches, specifically in the BLA of 3-month-old 3xTg-AD mice compared to WT. These observations indicate that these cells undergo a shift toward a less–ramified morphology, in line with proliferative microglial phenotypes (82) and potentials pro-inflammatory transcriptomic changes (83).

Our data also revealed a significant increase in the expression of TJ occludin and ZO-1 proteins in the HPC and the BLA of 3-month-old 3xTg-AD mice, associated with intact BBB permeability to FITC-dextran of 40 kDa, indicating that permeability to large molecules is unchanged. This result therefore demonstrates that TJs are not altered in 3-month-old 3xTg-AD mice, which is consistent with the concept of preserved BBB in AD mouse models (84). Consistent with our previous data, reports demonstrated the absence of BBB opening in 3xTg-AD mice aged 6 to 18 months (20,29). However, our previous studies demonstrated decreased BBB permeability to small molecules (sucrose) in the HPC of 6-month-old 3xTg-AD mice, and subsequently throughout the brain (20,29), as well as in the AD p25 model (28), suggesting a more restrictive BBB. Taken together, these results indicate that the early overexpression of TJ proteins observed in 3-month-old mice may reflect a BBB that is more restrictive to small molecules. It is noteworthy that these results can be put into perspective with the work of Shen and collaborators, who showed that the use of focused ultrasound with microbubbles in 8-month-old 3xTg-AD mice resulted in a localized opening of the BBB, associated with a reduction of cerebral deposits of Aβ and Tau, and an improvement in cognitive deficits (32). This suggests a link between BBB permeability and the pathological state, with barrier opening being potentially beneficial, whereas excessive closure might be detrimental. From this perspective, the early BBB more restrictive property we observed at 3 months may represent a mirror image of later modifications, supporting the idea that dynamic regulation of BBB integrity is an early and potentially modulatory step in the pathophysiology of AD in 3xTg-AD mice.

We performed a PCA including behavioral parameters and molecular markers assessed in 3-month-old 3xTg-AD and WT mice. PC1 clearly separated 3xTg-AD from WT mice, with strong contributions from anxious- and depressive-like behaviors, deficits in cognitive flexibility, and molecular markers including increased ZO-1 expression in the DG and the CA1, as well as microglial density in the DG and average length of microglial branches in the BLA. The regional variation highlights that early molecular changes in BBB TJ and glial cells closely associate with the emergence of BPSD-like behaviors in 3xTg-AD mice, suggesting a potential mechanistic link between barrier function, neuroimmune changes and behavioral dysregulation at this early stage of pathology.

Interestingly, several studies have established a link between anxious- and depressive-like behaviors and alterations in adult neurogenesis within DG. Notably, boosting neurogenesis in 3xTg-AD mice is sufficient to prevent abnormal behaviors as increases time spent in the center of the OF (76). DG neurogenesis has also been associated with cognitive flexibility (85), a modality we assessed here using an operant RL task. It has been proposed that DG neurogenesis partially relies on the neurovascular system (86) and that a finely tuned BBB permeability is necessary to support proper neurogenesis (87). In this context, the early BBB reinforcement we observed in 3-month-old 3xTg-AD mice, together with regional microglial alterations, could potentially impact adult DG neurogenesis, contributing to the emergence of BPSD-like behaviors and cognitive rigidity. Indeed, it has been proposed that an impaired adult hippocampal neurogenesis could provide a link between depression and AD (88), and a review highlighted the contribution of this neurogenesis both to mood and cognitive flexibility (85). This perspective provides a mechanistic framework linking early BBB and glial changes to behavioral and cognitive deficits in AD and suggests that modulation of the NGVU could be an important factor in the progression of the pathology.

In conclusion, our study demonstrates that 3-month-old 3xTg-AD mice already exhibit early BBB restriction and regional microglial alterations in the HPC and BLA, associated with anxiety- and depression-like behaviors as well as deficits in cognitive flexibility. Importantly, these BPSD-like features emerge before significant amyloid or tau pathology, a stage not previously explored in detail. Our findings provide new evidence linking early NGVU and neuroinflammatory changes to behavioral dysregulation in AD, highlighting the NGVU as a potential target for early intervention.

## Acknowledgements

The authors thank the Animal Facilities of Besançon for technical support, PhD Pierre-Yves Risold for his valuable assistance with the use of the microscope during the immunofluorescence experiments, Christophe Houdayer and Bahrie Ramadan for their technical support.

## Declarations

### Ethics approval

All animal procedures were conducted in accordance with the Guide for the Care and Use of Laboratory Animals (NIH), the Animal Research: Reporting of In Vivo Experiments (ARRIVE) guidelines, and the European Union regulations on animal research (Directive 2010/63/EU). They were approved by the University of Marie and Louis Pasteur Animal Care and Use Committee (CEBEA-58).

### Consent for publication

Not applicable

### Data availability

Data will be shared under reasonable request.

### Authors’ contributions

FB, AE, LC and GBC conceived and designed the study. GBC and TN performed the behavioral experiments. GBC, AE, FB and TN analysed the behavioral data. FB, AE, GBC and DM were involved in tissue collection. GBC and DM performed and analysed the FITC-dextran permeability assay. GBC, CM, SDC, FB and AE performed immunofluorescence and/or analysed the related data. GBC performed the western blot experiments. GBC, FB, AE and LC were involved in the statistical analysis and wrote the original draft of the manuscript. GBC and FB created the figures. All authors reviewed and edited the manuscript and approved the final version. FB and AE supervised the study and provided funding, resources and direction.

